## Supplementary material for "Circulating Small Extracellular Vesicle RNA Profiling for the Detection of T1a stage Colorectal Cancer and Precancerous Advanced Adenoma": Suppl figures.docx

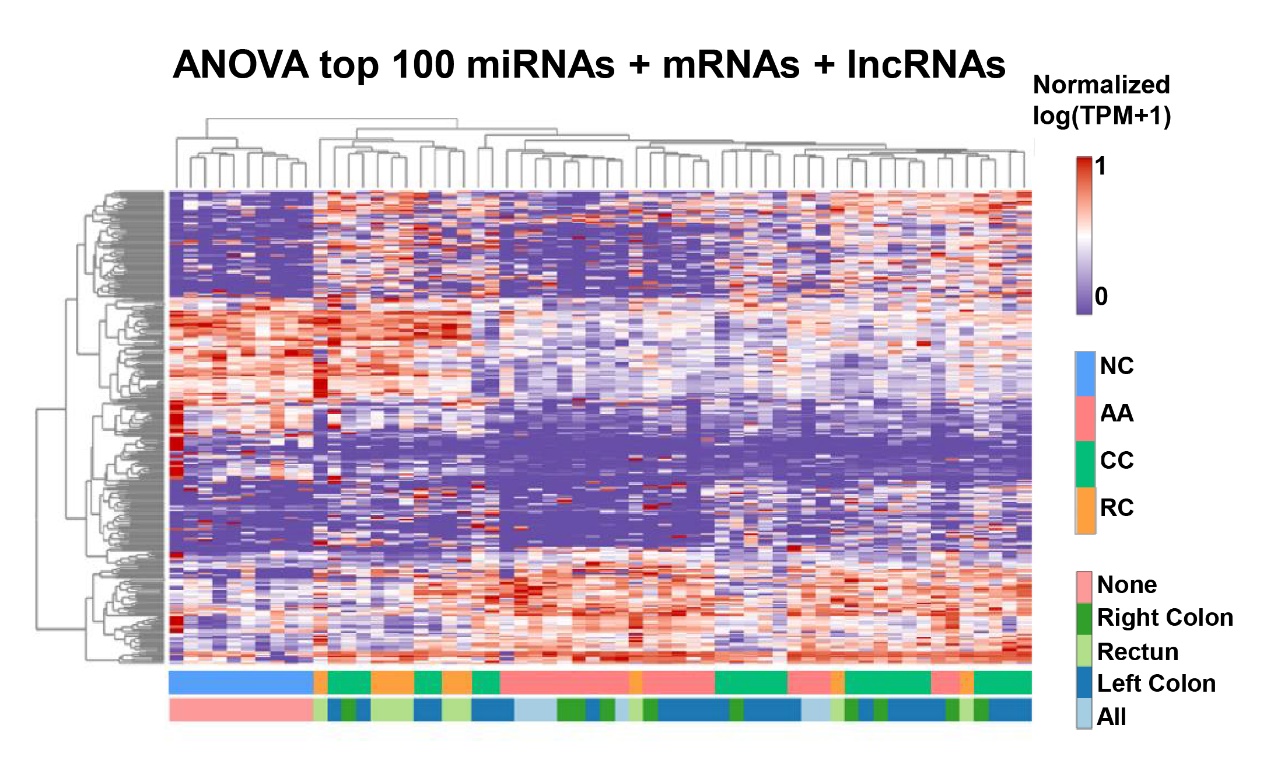

Figure 2—figure supplement 1.

The hierarchical clustering results of Top 100 miRNAs/mRNAs/lncRNAs.

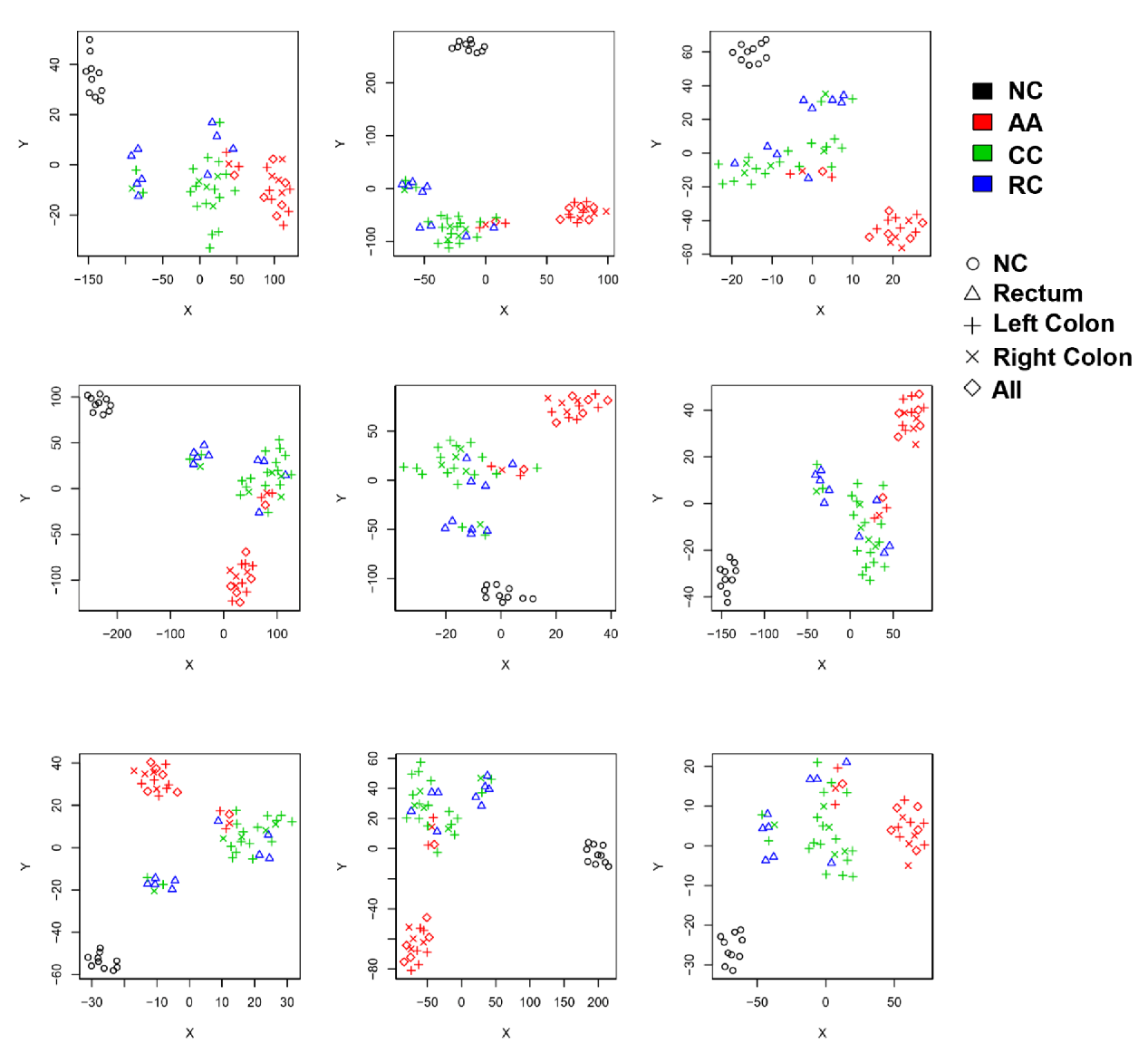

Figure 2—figure supplement 2.

Unsupervised t-SNE clustering by those 200 RNAs with 9 repeats.

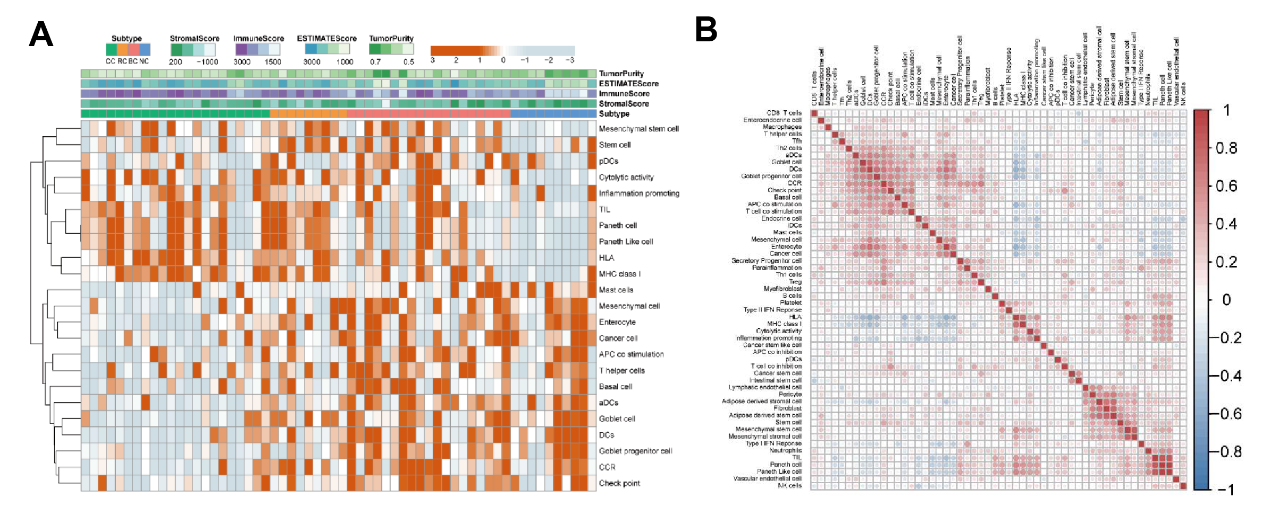

Figure 3—figure supplement 1.

(A) The hierarchical clustering heatmap of different cell features in all sEV samples. (B) Correlation among all cell-specific features.

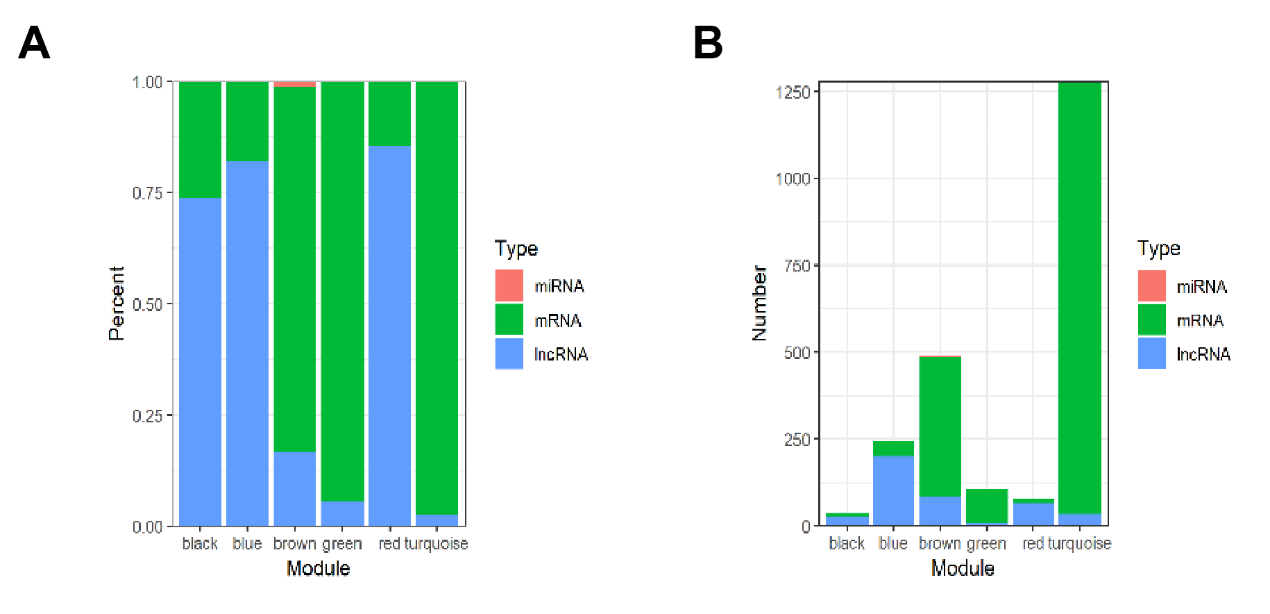

Figure 4—figure supplement 1.

(A) Percentage barplot of the RNA composition of different modules (only DEGs with kME>0.7).

(B) Barplot of module composition of different modules (only DEGs with kME>0.7).

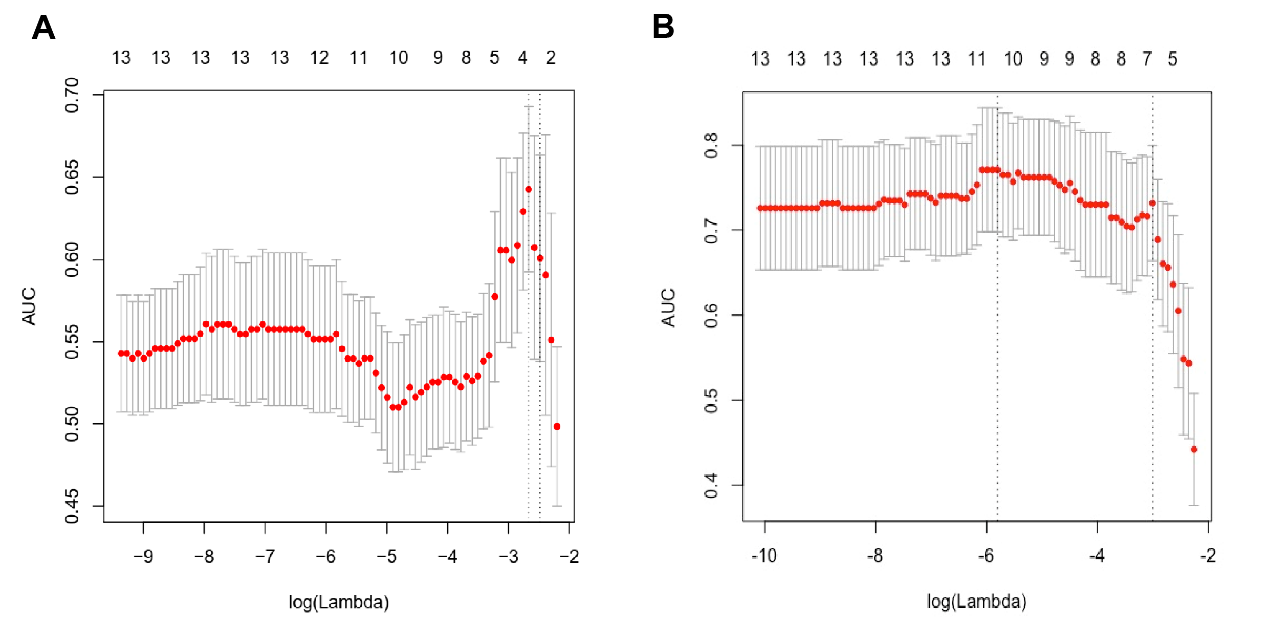

Figure 7—figure supplement 1.

(A) Performance of Lasso regression in variable selection to identify CRC.

(B) Performance of Lasso regression in variable selection to identify AA.

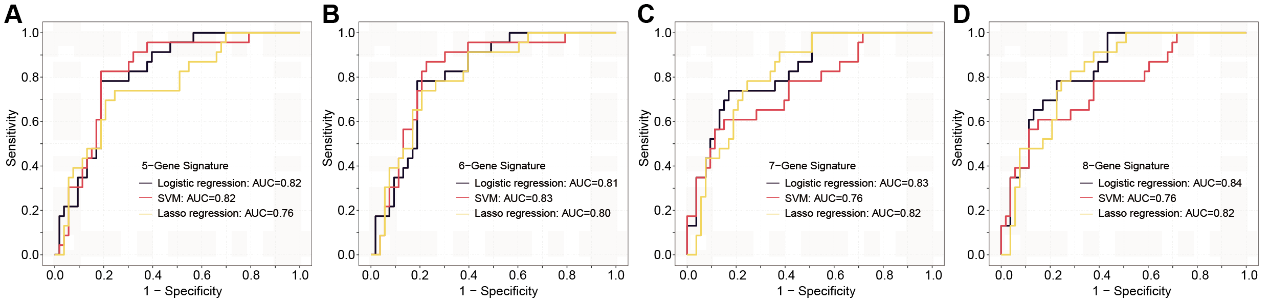

Figure 7—figure supplement 2.

The ROC analysis of different sEV-RNA signatures in the prediction of Stage I CRC patients by different algorithms (a: 6-gene panel; b: 7-gene panel; c: 8-gene panel; d: 9-gene panel).

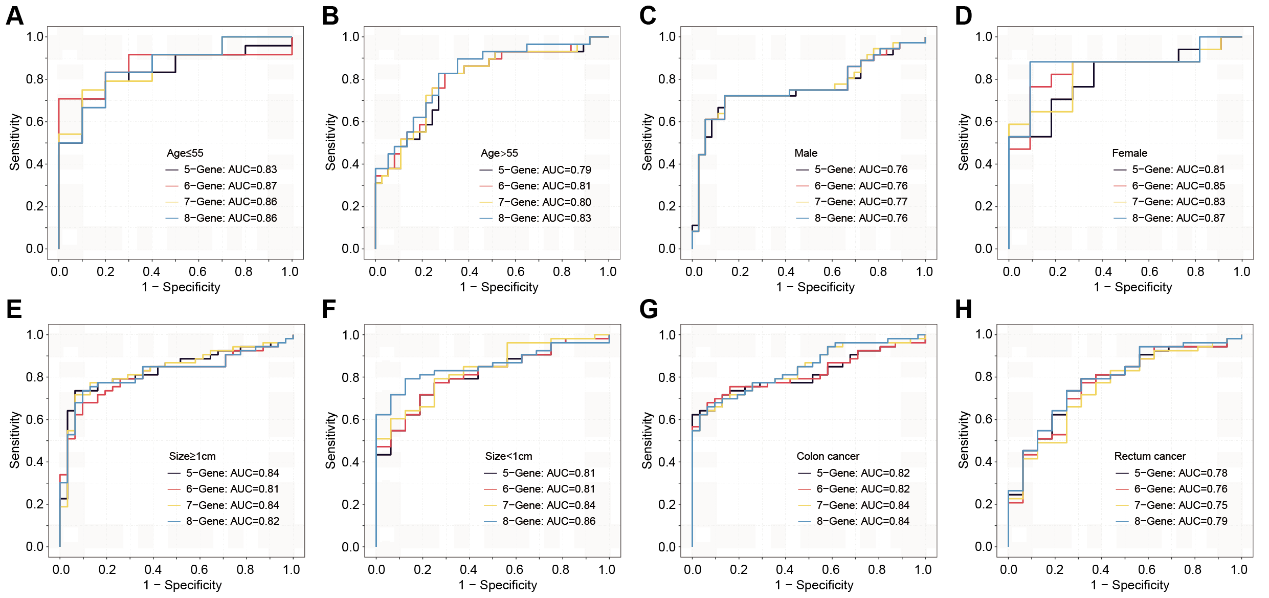

Figure 7—figure supplement 3.

The ROC analysis of different sEV-RNA signatures for predicting CRC patients using the Lasso regression algorithm in different clinical parameters (ab: age; cd: gender; ef: tumor size; gh: anatomical position).
