## Supplementary material for "Circulating Small Extracellular Vesicle RNA Profiling for the Detection of T1a stage Colorectal Cancer and Precancerous Advanced Adenoma": Suppl methods.docx

**Supplementary methods**

**PART1.** **Morphological characterization of sEVs**

**Nanoparticle tracking analysis (NTA)**

sEVs suspension (ranging from 1×10^7^/mL to 1×10^9^/mL) was detected by the ZetaView PMX 110 (Particle Metrix, Meerbusch, Germany) with a 405nm laser to determine the size and quantity of sEVs. A 60s video was taken with a 30 frames/s frame rate, and the movements of those nanoparticles were analyzed using ZetaView 8.02.28.

**Transmission electron microscopy (TEM)**

About 15µL sEVs suspension was placed on a copper mesh and incubated at room temperature for 10min. After washing with sterile distilled water, the sEVs solution was contrasted by the uranyl-acetate solution for 1min. Then the sample was dried for 2min and observed under a TEM microscope (JEOL-JEM1400, Tokyo, Japan).

**PART2. RNA sequencing, primary data processing and analysis**

**RNA library preparation and sequencing**

For long RNA libraries, a total amount of 5 ng RNA per sample was used as input material for sequencing libraries using the Ovation® SoLo RNA-Seq Library Preparation Kit (NuGEN, CA, USA) following the manufacturer’s recommendations and index codes were added to attribute sequences to each sample. For small RNA libraries, 2.5 ng RNA per sample was used as input material for the RNA sample preparation. Sequencing libraries were generated using NEB Next Multiplex Small RNA Library Prep Set for Illumina (NEB, USA) following the manufacturer’s recommendations and index codes were added to attribute sequences to each sample.

Then PCR products were purified (AMPure XP system) and library quality was assessed on the Agilent Bioanalyzer 2100. The clustering of the index-coded samples was performed on a cBot Cluster Generation System using TruSeq PE Cluster Kitv3-cBot-HS (Illumia). After cluster generation, the library preparations were sequenced on an Illumina Hiseq platform and paired-end reads were generated.

**Identification, quantification, and differential expression analysis of miRNAs**

The clean reads were aligned to the reference GRCh38 using Bowtie tools. Annotated by the Silva database, GtRNAdb database, Rfam database, and Repbase, ribosomal RNA (rRNA), transfer RNA (tRNA), small nuclear RNA (snRNA), small nucleolar RNA (snoRNA), other ncRNA and repeats were filtered before further analysis. The remaining reads were used to quantify known miRNA and predict new miRNA by compared to miRNAs from miRbase and GRCh38, respectively. Read count for each miRNA was obtained from the mapping results, and TPM was calculated. TMM normalization was performed and DEGs analysis of any two groups was conducted using the Mann Whitney U test with P-value < 0.05, and Fold change > 1.5.

**Identification of mRNAs and lncRNAs**

Raw data (raw reads) of fastq format were firstly processed through in-house Perl scripts. In this step, clean data (clean reads) were obtained by removing reads containing adapter, reads containing ploy-N, and low-quality reads. At the same time, Q20, Q30, GC-content, and sequence duplication level of the clean data were calculated. All the downstream analyses were based on clean data with high quality. Paired-end clean reads were aligned to the reference GRCh38 using TopHat2/Bowtie2. Mapped reads were used for the quantification of mRNA level and differential expression analysis.

For lncRNA analysis, the transcriptome was assembled using the Cufflinks and Scripture based on the reads mapped to the reference genome. The assembled transcripts were annotated using the Cuffcompare program from the Cufflinks package. The unknown transcripts were used to screen for putative lncRNAs. Three computational approaches including CPC (Coding Potential Calculator)/CNCI (Coding-Non-Coding Index)/Pfam were combined to sort non-protein coding RNA candidates from putative protein-coding RNAs in the unknown transcripts. CPC is a sequence alignment-based tool used to assess protein-coding capacity. By aligning transcripts with known protein databases, CPC evaluates the biological sequence characteristics of each coding frame of the transcript to determine its coding potential and identify non-coding RNAs.^1^ CNCI analysis is a method used to distinguish between coding and non-coding transcripts based on adjacent nucleotide triplets. This tool does not rely on known annotation files and can effectively predict incomplete transcripts and antisense transcript pairs.^2^ Pfam divides protein domains into different protein families and establishes statistical models for the amino acid sequences of each family through protein sequence alignment.^3^ Transcripts that can be aligned are considered to have a certain protein domain, indicating coding potential, while transcripts without alignment results are potential lncRNAs. Putative protein-coding RNAs were filtered out using a minimum length and exon number threshold. Transcripts above 200 nt with more than two exons were selected as lncRNA candidates and further screened by CPC/CNCI/Pfam. We distinguished lncRNAs from protein-coding genes by intersecting the results of the three determination methods mentioned above. Considering the current limited understanding of various sEV-lncRNAs in research, further exploration is necessary in the future to better elucidate the roles of different RNA categories involved in the tumorigenesis process.

**Quantification and differential expression analysis of mRNAs and lncRNAs**

Stringtie was used to calculate FPKMs of coding genes in each sample. Gene FPKMs were computed by summing the FPKMs of its all alternatively spliced transcripts. Genes with median FPKM<5 were regarded as low abundance genes and excluded in the subsequent analysis. TMM normalization was performed and differentially expressed genes (DEGs) analysis of any two groups was conducted using the Mann Whitney U test with cutoff P-value < 0.05, and Fold change > 1.5.

**Selection of Predictive Biomarkers**

To ensure the predictive performance of the sEV-RNA signature, candidate sEV-RNAs were ultimately selected based on their fold change in colorectal cancer/ precancerous advanced adenoma, absolute abundance, and module attribution. In detail, we initially selected the top 10 RNAs from each category (mRNA, miRNA, and lncRNA) with a fold change greater than 4. In cases where fewer than 10 RNAs were meeting this criterion, all RNAs with a fold change greater than 4 were included. Subsequently, we filtered out RNAs with low abundance, and we selected the top-ranked RNAs from each module based on the fold change ranking for inclusion in the final model.
