## Supplementary material for "Circulating Small Extracellular Vesicle RNA Profiling for the Detection of T1a stage Colorectal Cancer and Precancerous Advanced Adenoma": Suppl Table 12.docx

**Supplementary Table 12. Participants’ characteristics for the training and validation cohorts.**

|  | Training cohort (n=60) | | |  | Validation cohort (n=124) | | |
| --- | --- | --- | --- | --- | --- | --- | --- |
|  | CRC | AA | NC |  | CRC | AA | NC |
| Number | 31 | 19 | 10 |  | 47 | 24 | 53 |
| Sex |  |  |  |  |  |  |  |
| Male | 18 | 11 | 6 |  | 36 | 20 | 36 |
| Female | 13 | 8 | 4 |  | 11 | 4 | 17 |
| Age |  |  |  |  |  |  |  |
| >55 | 23 | 13 | 7 |  | 37 | 16 | 30 |
| ≤55 | 8 | 6 | 3 |  | 10 | 8 | 23 |
| Stage |  |  |  |  |  |  |  |
| I | 31 | - | - |  | 23 | - | - |
| II-III | 0 | - | - |  | 24 | - | - |
| Tumor size |  |  |  |  |  |  |  |
| ≥1cm | 25 | 10 | - |  | 31 | 14 | - |
| <1cm | 6 | 9 | - |  | 16 | 10 | - |
| Location |  |  |  |  |  |  |  |
| Right colon | 5 | 5 | - |  | 9 | 5 | - |
| Left colon | 17 | 8 | - |  | 22 | 13 | - |
| Rectum | 9 | 6 | - |  | 16 | 6 | - |
| Yamada_subtype | |  |  |  |  |  |  |
| I | 8 | 7 | - |  | 10 | 7 | - |
| II | 8 | 6 | - |  | 9 | 9 | - |
| III/IV | 15 | 6 | - |  | 28 | 8 | - |
