## Supplementary material for "Circulating Small Extracellular Vesicle RNA Profiling for the Detection of T1a stage Colorectal Cancer and Precancerous Advanced Adenoma": Suppl Table 13.docx

**Supplementary Table 13. Transcripts and sequence of their primers and** [**probe**](javascript:;)**s.**

| RNA | primer | sequence | Length (bp) |
| --- | --- | --- | --- |
| *miR-3615* | RT-primer | GTCGTATCCAGTGCAGGGTCCGAGGTATTCGCACTGGATACGACGAGCCG | 21 |
|  | Forward | CCTCTCTCGGCTCCTCG |  |
|  | Probe | TCGCACTGGATACGACGAGCCG |  |
| *miR-425-5p* | RT-primer | GTCGTATCCAGTGCAGGGTCCGAGGTATTCGCACTGGATACGACTCAACG | 23 |
|  | Forward | CGCAATGACACGATCACTCC |  |
|  | Probe | TTCGCACTGGATACGACTCAACG |  |
| *miR-106b-3p* | RT-primer | GTCGTATCCAGTGCAGGGTCCGAGGTATTCGCACTGGATACGACGCTGC | 22 |
|  | Forward | CACCGCACTGTGGGTACT |  |
|  | Probe | ATTCGCACTGGATACGACGCTGC |  |
| *Let-7f-5p* | RT-primer | GTCGTATCCAGTGCAGGGTCCGAGGTATTCGCACTGGATACGACGGAAAG | 22 |
|  | Forward | AGCGCCTATACAGTCTACTGT |  |
|  | Probe | TCGCACTGGATACGACGGAAAGAC |  |
| *miR-320a-3p* | RT-primer | GTCGTATCCAGTGCAGGGTCCGAGGTATTCGCACTGGATACGACGGAAAG | 22 |
|  | Forward | AGCGCCTATACAGTCTACTGT |  |
|  | Probe | TCGCACTGGATACGACGGAAAGAC |  |
| *C19orf43* | Forward | GCAGAAGGAAACCGAGATGC | 155 |
|  | Reverse | GAGGAAGACAGAAAGAGCCAGAG |  |
|  | Probe | TCCCGCAGCCGTGGACGATTCT |  |
| *TOP1* | Forward | CTGTAGCCCTGTACTTCATCGAC | 150 |
|  | Reverse | TCTACCACATATTCCTGACCATCC |  |
|  | Probe | CCTTCCTCCTTTTCATTGCCTGCTCT |  |
| *PPDPF* | Forward | CGCCTGGGTTCCACTTCCA | 171 |
|  | Reverse | CTCTGCGGACTCCAACACC |  |
|  | Probe | AAAAGAAGCTGGCCCACCAATGACCC |  |
| *MT-ND2* | Forward | GTATTTCCTCACGCAAGCAACC | 144 |
|  | Reverse | CTTGGGTAACCTCTGGGACTC |  |
|  | Probe | ACCGCATCCATA+AT+C+CTT+CTAATAGC |  |
| *HIST2H2AA4* | Forward | CATCATCCCTCGTCACCTCCA | 160 |
|  | Reverse | TCACTTGCCCTTTGCCTTGTG |  |
|  | Probe | CCATCCGCAACGACGAGGAACTGAAC |  |
| *lnc-MSI1-2:1* | Forward | TGCCAATGCCGACTATATTTCAAG | 110 |
|  | Reverse | GCAGTATCGTAGCCAATGAGGTTT |  |
|  | Probe | CTTTTCCCAATACCCCGCCATGACGA |  |
| *lnc-MKRN2-42:1* | Forward | TCCGCACTAAGTTCGGCATC | 107 |
|  | Reverse | TTTTGACCTGCTCTGTTTCCG |  |
|  | Probe | TCCTTAGGCAACCTGGTGGTCCC |  |
| *LNC-EV-9572* | Forward | ATCCACCTGCCTCAGCCTTC | 117 |
|  | Reverse | TCCCAAGATGGAATCAGGAAGAG |  |
|  | Probe | TCACTGAGGCTGACCTGGCAAACTT |  |

+ suggested a location modified with locked nucleic acid (LNA)
